# Tissue determinants of the human T cell receptor repertoire

**DOI:** 10.1101/2024.08.17.608295

**Authors:** Suhas Sureshchandra, James Henderson, Elizabeth Levendosky, Sankalan Bhattacharyya, Jenna M Kastenschmidt, Andrew M Sorn, Mahina Tabassum Mitul, Aviv Benchorin, Kyle Batucal, Allyssa Daugherty, Samuel JH Murphy, Chandrani Thakur, Douglas Trask, Gurpreet Ahuja, Qiu Zhong, Annie Moisan, Andreas Tiffeau-Mayer, Naresha Saligrama, Lisa E Wagar

## Abstract

98% of T cells reside in tissues, yet nearly all human T cell analyses are performed from peripheral blood. We single-cell sequenced 5.7 million T cells from ten donors’ autologous blood and tonsils and sought to answer key questions about T cell receptor biology previously unanswerable by smaller-scale experiments. We identified distinct clonal expansions and distributions in blood compared to tonsils, with surprisingly low (1-7%) clonal sharing. These few shared clones exhibited divergent phenotypes across bodily sites. Analysis of antigen-specific CD8 T cells revealed location as a main determinant of frequency, phenotype, and immunodominance. Finally, diversity estimates from the tissue recalibrates current repertoire diversity estimates, and we provide a refined estimate of whole-body repertoire. Given the tissue-restricted nature of T cell phenotypes, functions, differentiation, and clonality revealed by this dataset, we conclude that tissue analyses are crucial for accurate repertoire analysis and monitoring changes after perturbing therapies.

## INTRODUCTION

T cells are fundamental players in the adaptive immune system that can recognize a remarkable diversity of peptide antigens through their T cell receptor (TCR) repertoire^1,2^. While blood samples are often used to monitor changes in the human TCR repertoire, they represent less than 2% of the body’s T cells^3^. Most T cells reside in the lymphatics (spleen, lymph nodes, tonsils) and non-lymphoid tissues (gut, skin, lung, etc.), where they surveil for infections and maintain tissue homeostasis^4–7^. Specialized T cell subsets serve unique tissue-based functions^8^. For example, tissue-resident memory T cells (TRM) are crucial for local immune responses^9,10^ and homeostasis^11^, and T follicular helper cells (TFH) are essential for germinal centers^12–14^. These specialized T cells are often absent or rarely detected in circulation, so tissue-based analyses are essential to understand the full scope of T cell diversity and function^15^. The extent of phenotypic mixing and repertoire compartmentalization between blood and tissue T cells remains unclear, largely due to low-scale sampling. It is likely that T cell location influences its phenotype and TCR specificity. Current evidence suggests that while T cell clones with common ancestors exhibit similar transcriptional profiles, single naive clones can differentiate into various phenotypes during infection^16–19^ or tumor progression^20^. Although much progress has been made over the last decade, several critical questions remain unanswered. First, what are the patterns of phenotypic concordance or divergence within clones shared between blood and tissue? Second, since tissue sites maintain T cell clones that respond to localized cues, is the relationship between a cell’s TCR and phenotype dependent on its location? Finally, is the extent of clonal sharing dictated by the nature of the antigen itself?

TCR repertoire diversity is also shaped by various host factors^21^ and environmental factors. Age is a major determinant of TCR diversity^22–25^ and serves as a proxy for both changes in T cell development and accumulation of immune memory, encompassing exposure to pathogens and vaccines, self-reactivity, and the cumulative impact of chronic diseases^26^. Importantly, memory T cells increase significantly in the first decade of life^27,28^, with early antigen exposures shaping TCR diversity, which is later refined by clonal selection^29^. A decline in T cell function and diversity is associated with increased susceptibility to disease^30^. An accurate estimation of TCR diversity in humans is, therefore, important for understanding effective control of infections and subsequent protection^31,32^.

Current estimates of peripheral TCR diversity have relied heavily on single (beta) chain sequencing of up to a million T cells to extrapolate overall bodily diversity^23,33,34^. While a broad range of T cells (>10^20^) is theoretically possible, our body has only ∼4.5 × 10^11^ T cells^3^, with actual TCR diversity significantly lower due to thymic selection and clonal expansion following antigen-specific response^35^. On the other hand, a single TCR can recognize a variety of peptide-MHC complexes^36–38^, reducing the diversity needed to recognize any antigen^39–41^. Estimates of T cell clonotypes in the human repertoire range from 3 × 10^6^ (β chain diversity)^42^ to 10^7^ - 10^8^ ^23,33,43^. A major challenge in reconciling these estimates is under sampling, as statistical estimators struggle with poor sampling, particularly for small clones and singletons^26,35^. However, recent improvements in single-cell sequencing offer new opportunities for profiling paired TCR chains and T cell transcriptomes from hundreds of thousands of single cells in an unbiased manner.

In this study, we collected paired gene expression and TCR data from a cohort of ten donors, spanning from infancy to adulthood, and investigated both phenotypic and clonal segregation of T cells between blood and tissue. Tissue T cells were isolated from tonsils, which is a unique and accessible site for studying the influence of both lymphoid and mucosal tissue environments. By examining transcriptional and TCR sequencing data from 5.7 million T cells, we show limited clonal sharing between blood and tissue and that blood sampling alone significantly underestimates clonal and phenotypic T cell diversity in the human body. Our analyses also revealed differences in antigen-specific T cell frequencies, phenotypes, and immunodominance restricted by bodily site. These findings shed new light on our current understanding of TCR repertoire biology and have significant implications for a new era of tissue-focused vaccine and immunotherapy strategies.

## RESULTS

### Comprehensive analysis of phenotypic and clonal diversity of T cells in blood and tonsils

T cells in tissues acquire unique phenotypes and functional attributes compared to their counterparts in blood^44,45^. We posited that this compartmentalization of barrier sites from circulation would result in unique phenotypic and clonal characteristics reflecting tissue site-specific exposures. To date, many studies have analyzed and provided approximates of the complexity of human TCR repertoire^33,43^. However, technological limitations (single chain profiling of a couple of thousands of cells) and challenges in sampling (several donors, but not enough sampling depth, and lack of tissues) have, thus far, limited us from fully reconciling previous assumptions and estimates of human TCR repertoire^26^. Here, we seek to address these questions by deeply characterizing paired transcriptional and T cell receptor (TCR) repertoire data from Lβ+ T cells from both blood and tonsils **(Figure S1A)** from 10 donors, averaging about 300,000 single T cells per donor per tissue compartment (**Figure 1A**). Our primary cohort included demographically diverse donors with different exposure histories to estimate immune variation among individuals **(Table S1)**. Selected donors ranged in age from infancy to adulthood, and samples from both male and female participants were included. Donors were diverse in their prior exposure to herpesviruses, known to influence the TCR repertoire^46^, and their indication for surgery **(Figure 1A)**.

**Figure 1:**
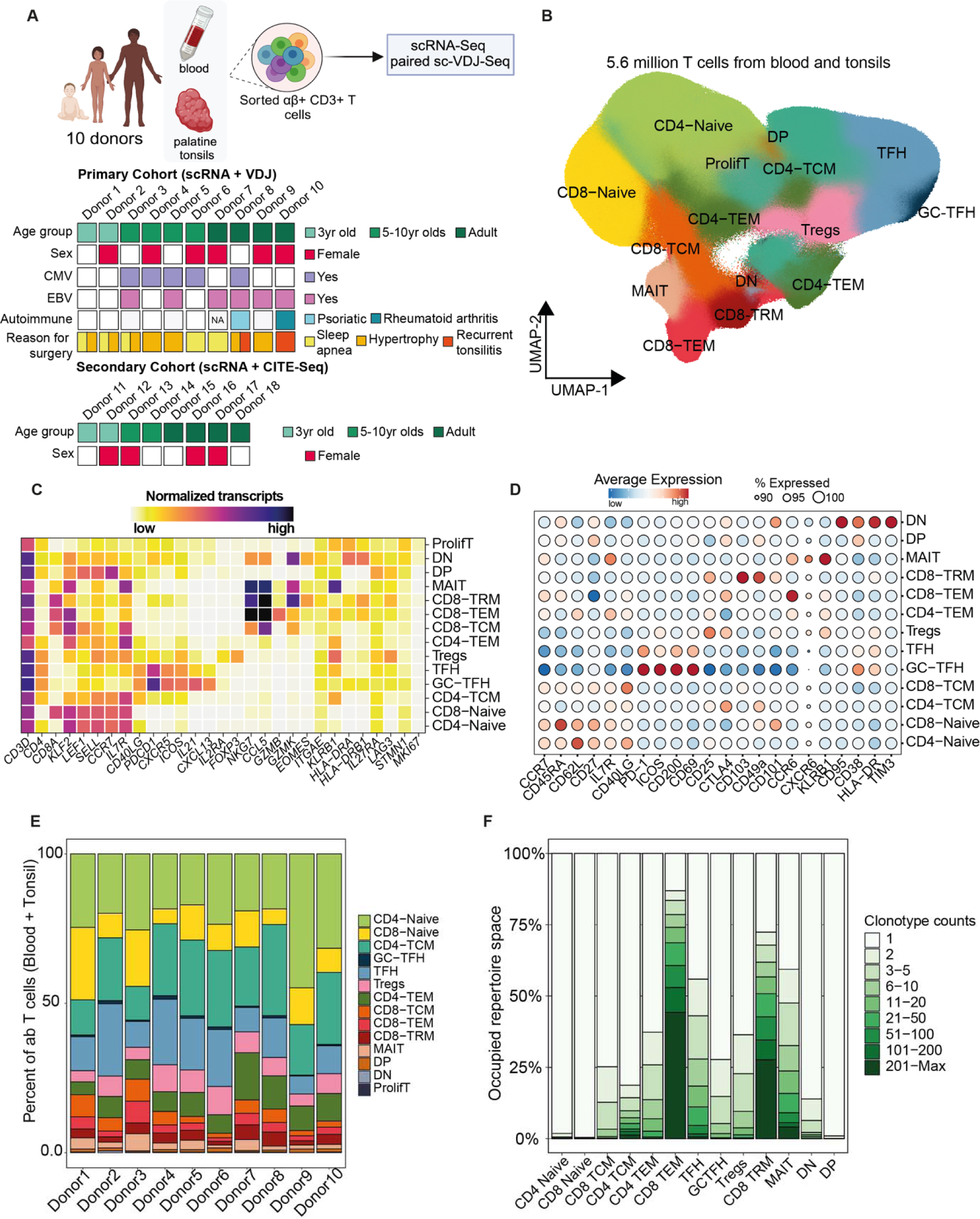
A high-resolution map of blood and tonsillar T cell subsets. (A) Experimental design and brief description of the primary and secondary cohort of human donors used in this study. (B) UMAP of 5.7 million cells from blood and tonsils. Level 3 cluster annotations are highlighted. (C) Heatmap of top distinguishing gene markers for each annotated cluster. (D) Bubble plot highlighting the relative expression of a subset of key cell surface markers from the secondary cohort of donors across L3 clusters. The bubble size indicates the percentage of cells in each cluster with detectable protein expression, and color indicates the magnitude of expression ranging from low (blue) to high (red). (E) Stacked bar graph comparing proportional breakdown of L3 clusters from each of the ten donors from the primary cohort. (F) Stacked bar graph comparing clone counts across T cell clusters from all donors. Only cells with productive TCRs were included in downstream analyses.

Data from blood and tonsils from each donor were integrated and clustered to identify and eliminate any contaminating non-Lβ+ T cells. Single cell profiles from the 10 donors were then integrated, normalized, and harmonized, resulting in a combined 5,728,381 cells (**Figure 1B; Figure S1B**). We integrated this dataset with blood and tonsillar Lβ+ T cells (59,859 cells) from a secondary cohort of 8 donors, where both gene expression and surface protein readouts of 32 T cell markers **(Table S2)** were captured using multimodal immune profiling. Cells were clustered by gene expression at a high resolution to capture rare cell states, resulting in 43 clusters. We carefully stratified these clusters into three levels of annotations based on both gene and protein markers, starting from a broad level 1 annotation (L1: CD4, CD8, double positive (DP), double negative (DN)) to level 2 annotation (L2: naive vs. memory subsets), and a refined level 3 annotation (L3: central memory (TCM), effector memory (TEM), regulatory T cells (Tregs), T follicular helper cells (TFH), etc. as represented by individual clusters) (**Figure 1B**). Numerous expected T cell phenotypic populations were identifiable based on a manual review of gene expression (**Figure 1C, Table S3**) and protein-level expression (**Figure 1D**) data.

All donors were well represented within each cluster (L3 annotation) (**Figure 1E**), although their proportions varied substantially between individuals. TFH subsets were more abundant in pediatric donors, whereas CD4 effector memory (TEM) frequencies increased with age (**Figure S1C**). Although the number of cells recovered from each donor was variable (ranging from 380,000 to 820,000), TCR sequence detection was efficient among all donors, indicating that each donor was well represented in the TCR repertoire analysis (**Figure S1D**). Overall, paired productive LβTCRs were recovered from roughly 3 million single cells. Productive TCRs were well represented in all major T cell clusters, with slightly lower recovery among mucosal-associated invariant T (MAIT) cells (**Figure S1E)**. As expected, naive T cell clusters were the most diverse, while memory CD8 T cells (effector memory and resident memory) and MAITs were highly clonal. TFH, which are largely restricted to lymphoid sites, were the most clonal subset within CD4 T cells **(Figure 1F)**. This high-resolution map of 5.7 million T cells is the largest (total and on a per-donor basis) dataset of immune cells profiled to date, encompassing multimodal readouts (transcriptomic, TCR repertoire, and protein-level expression) and will serve as a blueprint to compare TCR complexity in circulation and tissue.

### Compartmentalization and TCR sharing between tissue sites

To assess the extent of sharing vs. segregation of T cell phenotypes and clones between the circulation and tonsils, we stratified the dataset by their compartmental origin (blood vs. tonsil). As expected, naive CD4 and CD8 T cells were readily detected in both sites. However, T cells with certain phenotypes, such as TFH, CD8 resident memory (TRM), DP, and DN, were restricted primarily to tonsils, whereas CD8 effector and central memory cells were proportionally higher in circulation (**Figures 2A, S2A, and 2B**). Tregs were detected in both compartments, albeit at higher frequencies in the tonsils. In contrast, most MAIT cells were captured from blood rather than tonsils, as determined by both transcriptomic data and V/J gene usage.

**Figure 2:**
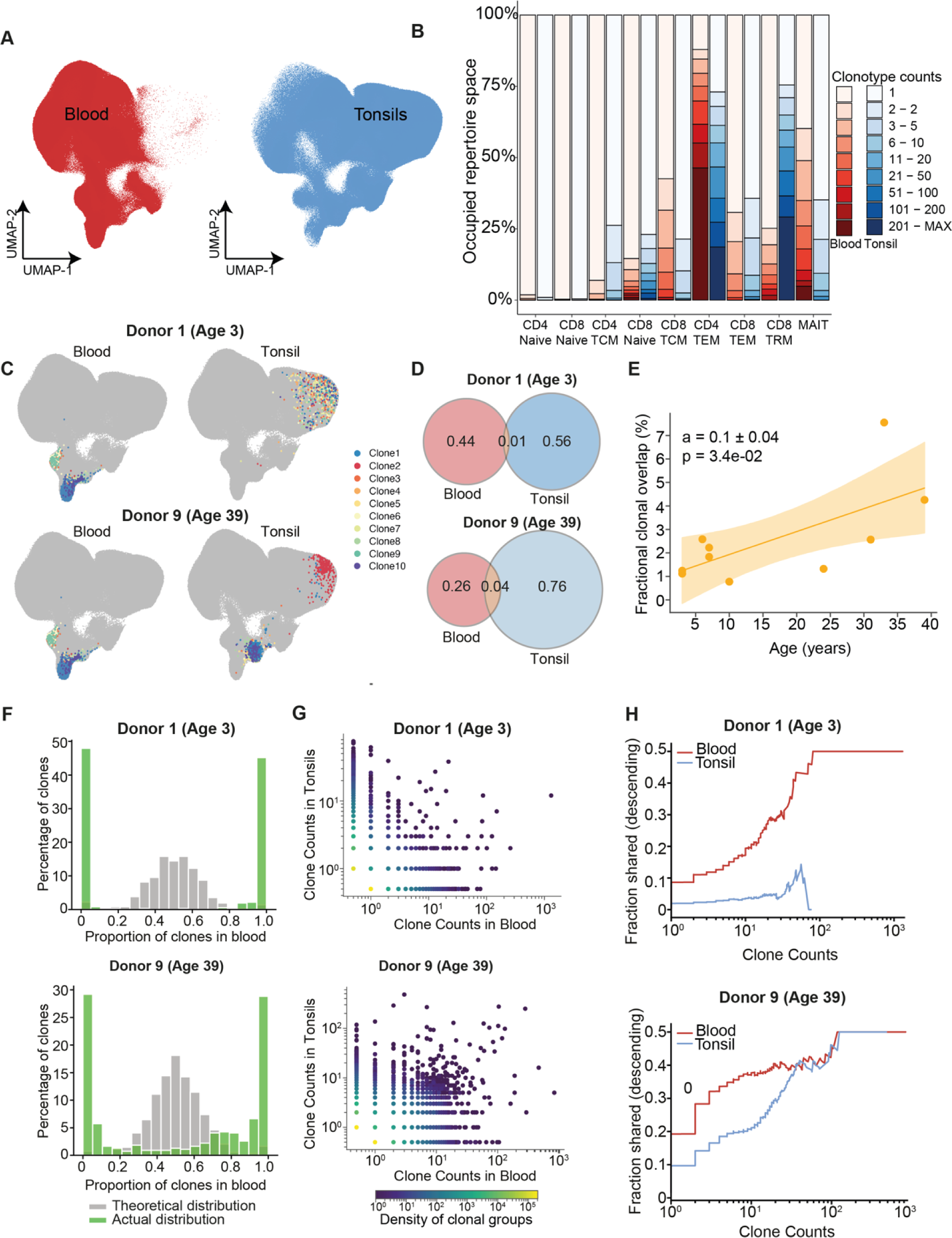
Compartmentalization of TCRs between tonsils and circulation. (A) UMAP of T cells from all donors split by tissue origin, i.e., blood or tonsil. (B) Stacked bar graph comparing the distribution of clone counts within each T cell cluster (L3 annotation) between blood and tonsils. (C) UMAP of T cells from blood (left) or tonsils (right) from two donors, Donor 1 and 9, with their top 10 clones from each compartment highlighted in color. (D) Venn diagrams comparing clonal overlap of T cells between blood and tonsils in the two representative donors. Values indicate the fraction of unique nucleotide sequences per donor. Overlap values indicate the number of unique nucleotide sequences present in both compartments as a fraction of the total number of unique sequences per donor. (E) Line graph comparing the fraction of clones (intersect values from D) shared between blood and tonsils in relation to donor age (represented on the x-axis) (F) Histograms of well-mixed (theoretical, grey) and true clonal sharing (green) of memory T cell compartments between blood and tonsils of the two representative donors. The x-axis represents the proportion of clonal frequencies in blood. (G) Scatter plots directly comparing clone counts in blood and tonsils from the two representative donors. Clones with members only in one compartment were given a clone count 0.5 in the other tissue. The color intensity of scatter points represents the density of clones within each compartment. (H) Clone count distribution for the two representative donors. y-axis represents the average fraction of clones of a particular size with a member in the second compartment. Clones are ranked by size, smallest to largest, from left to right.

We restricted downstream analysis to cells with productive TCRs, i.e., cells with at least one beta chain and one or two alpha chains. T cells with productive TCRs were well represented in both blood (ranging from 40-76%) and tonsils (68-83%) from each donor (**Figures S2C, S2D**). We examined the phenotypes of the largest clones in each compartment. While central memory (TCM) subsets were more clonal in tonsils (clonal proportions: 26% for CD4s, 23% for CD8s), TEMs (43% for CD4s and 88% for CD8s) and MAITs (60%) were more clonal in blood (**Figure 2B**). Consistent with the literature^47^, the largest clones in blood were CD8 TEM and MAIT cells (**Figures 2C, S3**). In contrast, the largest expanded clones in tonsils were of TFH phenotype in pediatric donors and mostly CD8 TRMs in adults (**Figures 2C, S3**), and the largest tonsil clones generally occupied more diverse transcriptional profiles than blood clones (**Figure 2C**). Since millions of paired TCR alpha and beta chain sequences were recovered, we assessed the proportion of T cell clones detected in both tissue sites. Unique clones were identified by exact nucleotide matching for alpha and beta TCR sequences, likely representing T cells from a common ancestor. Out of the total TCR repertoire, clonal overlap ranged from approximately 1-7% between blood and tonsils, depending on the donor (representative donors shown in **Figure 2D**). We established a clear correlation between clonal sharing across compartments and donor age (**Figures 2E, S4A**), with older donors showing more shared clones between tissue sites than younger donors. To determine whether the empirical clonal overlap reflected strong or weak sharing among bodily sites, we compared actual repertoire sharing to a theoretically well-mixed TCR distribution estimate. The estimate was generated by equalizing (via subsampling) the number of T cells in each tissue site for a given clone to account for clone size distribution biases. Overall, TCR sharing among tissues was heavily skewed from a theoretical perfect sharing (**Figures 2F and S4B**). Notably, many of the largest expanded T cell clones identified from tonsils were not detected in blood (**Figures 2G and S5A**), indicating that even highly expanded tonsil T cell clones are not always detectable by peripheral blood sampling. In contrast, most of the largest blood-derived T cell clones were identified in tonsils, though at much lower frequencies than in blood. This disparity was consistent across the clonal spectrum, with medium-sized clones from blood more likely to appear in tonsils (**Figures 2H and S5B**) than vice versa. Collectively, these findings suggest that T cell clonal expansions are often restricted to discrete bodily compartments and that blood represents a subsample of T cells from various sites.

### TCR diversity estimates from tonsils and blood

Current estimates of human TCR diversity were derived primarily from peripheral blood sampling and estimated to span several orders of magnitude, from 3 × 10^6^ (β chain diversity)^42^ to 10^7^ - 10^8^ ^23,33,43^, based on modeling of both alpha and beta chains. However, given the limited clonal sharing we observed between blood and tonsils, we aimed to determine a more accurate estimate of TCR diversity that covered the distribution of clones in both circulation and in tissue sites. A major bottleneck in diversity analysis of human TCR repertoire is undersampling^26^, coupled with the wide variability of clonal abundances. The distribution of clone counts within the repertoire is heavy-tailed^48^, with the proportion of clones exceeding clonal count thresholds decreasing with approximate power-law scaling (**Figure 3A**). While the largest clones comprised up to 10,000 cells (>1% of total cells), more than 90% of TCRs were represented by only a single cell in the sampled repertoire. Therefore, even at the scale of this dataset (about 20-fold higher per sample than a typical single-cell dataset for a single donor), rarefaction analysis showed no saturation of discovered clonotypes with increasing cell numbers (**Figure 3B**). Moreover, due to the broad distribution of clonal expansion sizes, we expect many expanded clones would not have been detected as such at the typically smaller dataset sizes of prior scTCRseq studies. Indeed, when subsampling cells for each donor, we observed that between 30-60% of memory clonotypes identified as singletons at the average sampling size of prior studies were part of expanded clonotypes when considered within the entire dataset (**Figure 3C**). These results underscore the challenges of insufficient sampling for accurate T cell diversity assessments.

**Figure 3:**
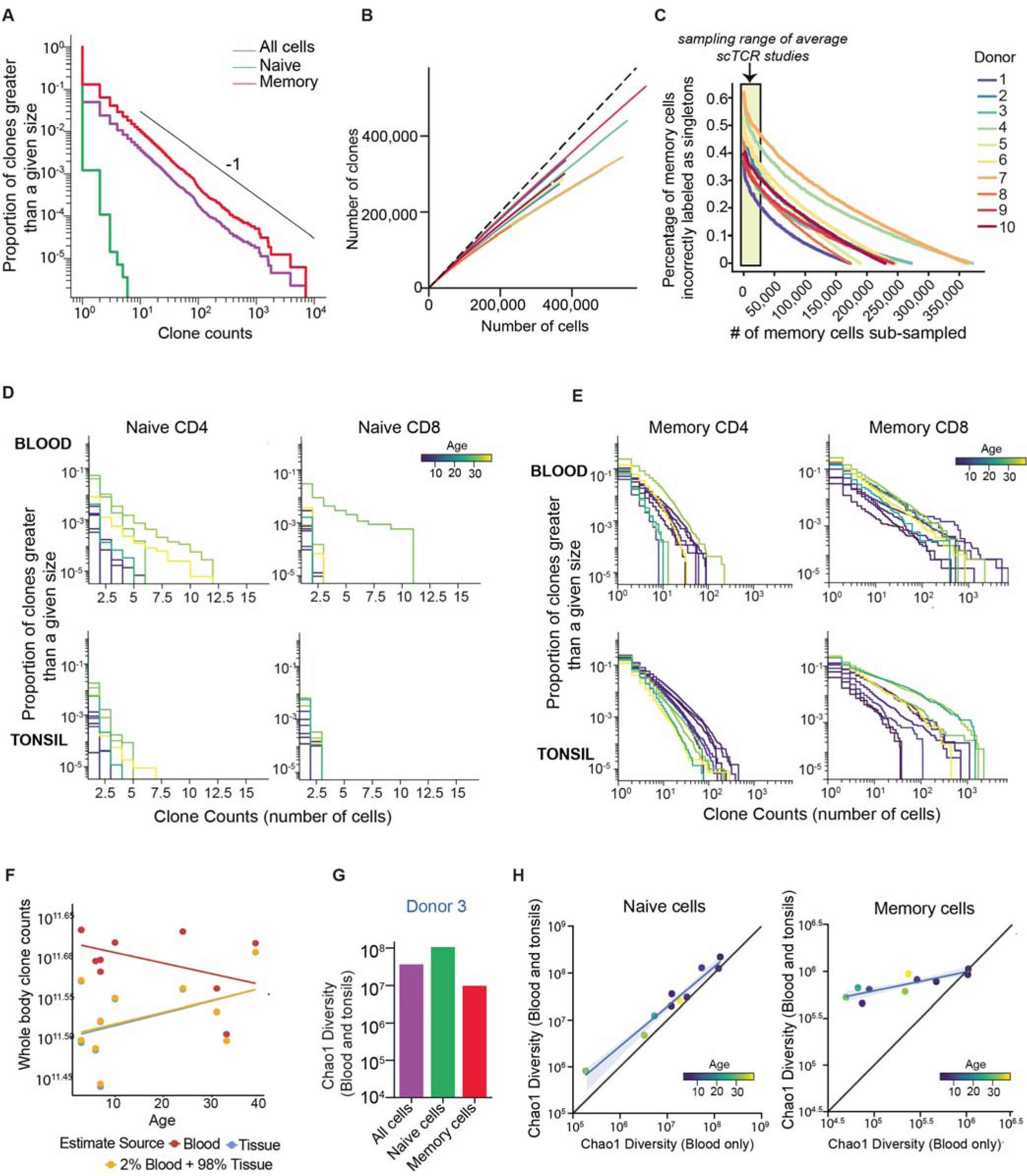
Diversity estimates of human blood and tissue TCR repertoire. (A) Clone count distribution of naive, memory, or total T cells for a representative donor, Donor 3. The y-axis represents the proportion of clones greater than a given size of the clone. (B) Rarefaction curves compare the number of clones relative to the number of cells sampled. Each line represents a donor, with blood and tonsillar T cells combined and singletons highlighted as a dashed line. (C) Comparison of fraction of memory T cell clones incorrectly assigned as singletons as a function of the numbers of cells sampled. The sampling size of a typical single-cell TCR dataset is highlighted in yellow. (D-E) Clone count distribution for (D) naive and (E) memory T cells from blood (top) or tonsils (bottom) for all ten donors (colored by age). The y-axis represents the proportion of clones greater than a given size of the clone. (F) Comparisons of whole-body TCR estimates from rarefaction analyses of each compartment extrapolated to total T cells in the human body. The x-axis represents the donor’s age. (G) Nonparametric (Chao1) estimates of TCR diversity for T cells (blood and tonsils combined) from Donor 3, split by their naive/memory phenotype. (H) Comparisons of Chao1 diversity of naive (left) or memory (right) T cells from blood and tonsils combined (y-axis) vs. blood cells alone (x-axis). Each dot represents a donor, and the color represents their age.

Next, we assessed the clone count distributions within CD4 and CD8 T cells, each separated by naive and memory phenotypes. We detected clonal expansions in all subsets (defined here by a minimal clone count of two cells), including within the naive CD4 and CD8 compartments (**Figures 3D, 3E**). Within naive CD4 T cells, as previously described^25,22,23^, these expansions were more frequent with increased donor age (**Figure 3D**) and were found independent of tissue site, albeit at a higher magnitude in blood. As expected, memory T cell populations contained significantly more clonally expanded TCRs (**Figures 3A, 3E**). Memory CD8 T cells were more clonal than memory CD4s, underscoring their divergent proliferative responses to antigens^49,50^. Differences in clonality were also observed based on tissue compartment, where memory CD4 (all donors) and memory CD8 (in adults) T cells were more clonal in tonsils compared to their blood counterparts (**Figure 3E**). Collectively, these findings highlight the role of homeostatic mechanisms, antigen exposure, and tissue-specific factors against the backdrop of aging in shaping the T cell repertoire.

Roughly 98% of T cells (of the 4.5 × 10^11^) in the human body are in lymphatics or non-lymphoid tissues^3^. We, therefore, asked if simple extrapolation of tonsils vs. blood repertoire rarefaction curves to whole body clone counts would predict different overall TCR richness estimates. Our analyses revealed a consistent trend of higher estimates from blood repertoire (average 3.95 × 10^11^ clones) vs. tonsils (average 3.35 × 10^11^ clones) (**Figure 3F**). However, differences in estimates were highly variable, ranging from 2-43% higher when using blood compared to tissue repertoire. To obtain statistically more robust lower bounds on TCR richness, we applied the nonparametric Chao151 estimator of species (i.e., clones) diversity to sampled T cell clone counts. Considering the differences in clonal distributions between naive and memory T cells, we analyzed Chao1 estimates not only for the entire repertoire but also separately for these subsets. Unexpectedly, we found that naive diversity estimates surpassed the diversity estimates from the total repertoire (representative donor in **Figure 3G**). These results indicate that applying diversity estimators based on T cell phenotype provides a more accurate estimate of total T cell diversity than considering the entire T cell pool. While memory T cells in the tonsils maintained similar diversity levels across donors, memory T cell diversity in the blood decreased with age (**Figure S6**). These findings suggest a preferential accumulation of expanded clones over time in the blood compared to tonsils, while naive T cell diversity declines with age in both compartments (**Figure S6**).

Finally, we investigated how relying solely on blood T cell data could influence diversity estimates compared to incorporating repertoire data from both blood and tissue. To this end, we compared Chao1 estimates for the naive and memory repertoire calculated using either all sampled cells from each donor or cells from blood alone (**Figure 3H**). While naive TCR diversity estimates were similar, memory diversity was underestimated by up to an order of magnitude in some donors when only blood T cells were sampled (**Figure 3H**).

### TCR-driven biases in T cell transcriptional state

Prior work has established that T cells differentiating from a common naive T cell are likely but not guaranteed to acquire similar differentiation profiles^16,17,19,52^. However, only a handful of studies have systematically examined the link between TCR sequence and T cell phenotype potential^53–55^. Consequently, we asked if there exists a predictable relationship between a cell’s TCR and its phenotype and whether that relationship is influenced by the cell’s presence in either blood or tissue. To address this question, we used the most detailed phenotypic annotation of each T cell cluster in the dataset, extending the annotations to Level 4 (L4) phenotypes (**Figure 4A**) based on the expression of specific costimulatory markers, activation states, and cytokines (**Figures 4B, S7A**, **and S7B**). Given the limited clone sharing between blood and tonsils, we first asked whether the few shared clones between tonsils and blood shared phenotypic features. Overall, we observed high concordance associated with CD4 vs. CD8 T cell lineage fate and naive vs. memory T cell differentiation status **(Figure 4C)**. As our phenotypic comparisons became more fine-grained (e.g., specific central/effector memory phenotypes and transcriptional profiles), concordance dropped for both blood and tonsil T cells (**Figure 4D**). CD8 memory subsets and MAITs exhibited high clonal sharing between blood and tonsils (**Figure 4E**). However, it was more probable that a clone found in both compartments acquired distinct transcriptional profiles based on its tissue site. The most common pairs with converging phenotypes in blood and tonsils were CD4 and CD8 TEMs, whereas the most common pairs with diverging phenotypes were clones with CD8 TRM phenotype in tonsil and CD8 TEM phenotype in blood, or CD4 TCM phenotype in tonsil and CD4 TEM phenotype in blood (**Figure 4F**).

**Figure 4:**
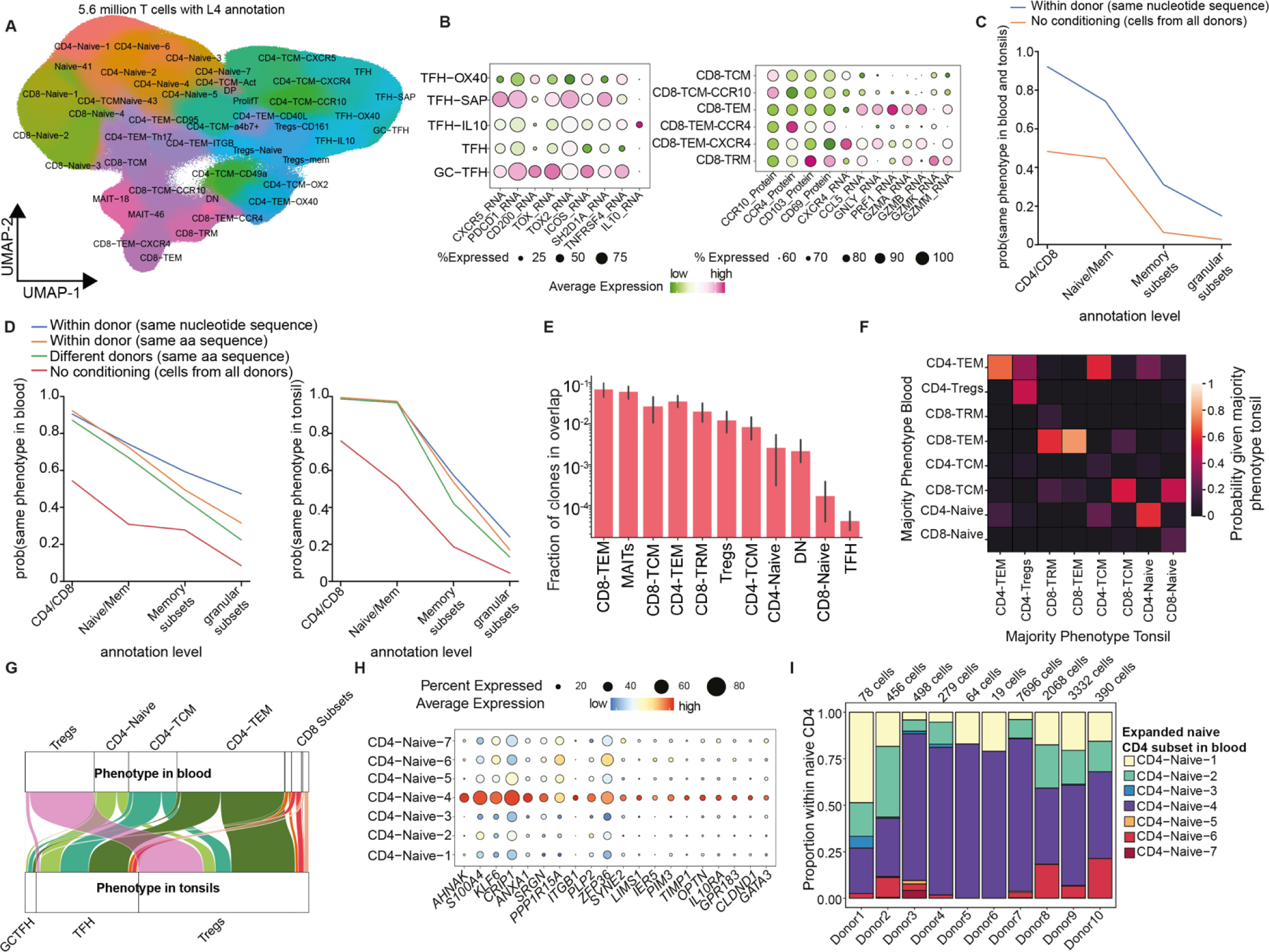
Relationship between TCR and T cell phenotype. (A) UMAP of 5.7 million T cells colored by annotation level 4 (L4). Cluster labels highlight L3 annotation in addition to highly expressed markers (tissue homing, cytokine, chemokine, costimulatory molecules, etc.). Bubble plots highlight distinct subsets of TFH (left) and memory CD8 T cells (right) subsets by differential RNA or surface protein expression. (C) Probability of a clone having identical phenotype between tissue compartments. Probabilities are compared across annotation levels (y-axis). A clone here is defined by identical nucleotide sequence matches. The probability of phenotypic sharing independent of the donor origin is highlighted in orange. (D) Compartment-specific probability (blood-left; tonsil-right) of two identical clones having identical phenotypes across annotation levels (x-axis). The blue line represents cells from the same donor with identical nucleotide sequences; orange represents cells from the same donor with identical amino acid sequences; green represents cells from all donors with identical amino acid sequences; red - cells from all donors with no conditioning (E) Bar graph highlighting the fraction of clones that have members in both blood and tonsils. Data are aggregated across donors. (F) Confusion matrix highlighting T cell phenotypes of shared clones in blood (y-axis) and tonsils (x-axis). Only clones shared between both compartments are highlighted here. Colors indicate the number of shared clones as a proportion of clones in tonsils. (G) Alluvial plot tracking blood phenotype of cells clonally related to tonsillar TFH, GC-TFH, and Tregs. (H) Bubble plot of top differentially expressed genes within naive CD4-4 subset relative to other naive CD4 subsets within L4 annotation. The size of the bubble represents the percent of cells within each cluster expressing the marker, whereas color represents normalized, scale transcript counts (I) Phenotype of expanded naive CD4 T cells (annotation level L4) across donors. Number of cells is highlighted.

To assess the extent of clonal sharing in different T cell phenotypes, we looked at subsets represented in both tonsils and blood. Tregs, for example, exhibited relatively high rates of clonal sharing between the sampled sites (**Figures 4E, 4F**)^56^. However, our analysis revealed non-Treg phenotypes in blood that were clonally related to tonsil Tregs, including CD4 TEM cells expressing tissue homing markers and a subset of naive CD4s (naive CD4-4) (**Figure 4G**). Conversely, T cell phenotypes restricted to tissue, such as TFH and GC-TFH, had matching TCR sequences to non-TFH cells in circulation, where they mainly manifested as TEM and TCM, expressed homing markers and, to some degree, were related to a naive cluster (CD4-4) **(Figure 4G**). Differential gene expression (DEG) analysis of this naive CD4-4 cluster relative to other naive CD4 clusters revealed up-regulation of homing (such as *ITGB1*) and activation (*KLF6* and *ANHAK*) markers (**Figure 4H**). Interestingly, this subset of naive CD4s accounted for most of the expanded naive CD4 T cells and was particularly enriched in adults (**Figure 4I**). Taken together, these results show not only a significant degree of differentiation plasticity for T cells, but also highlight the influence of location on phenotypic fate. We conclude that T cell phenotype in the blood does not reliably reflect the differentiation state and functional role of identical clones in tissue.

### Divergent differentiation fates of influenza-specific CD8 T cells derived from naive and memory pools following vaccination and infection

We next asked if memory CD8 T cell clones activated by pathogen or vaccine antigens maintain their phenotype or acquire distinct transcriptional states. Tonsil organoids (from 6 of the 10 analyzed donors) were stimulated with live-attenuated influenza vaccine (LAIV) or wild-type A/California/2009 H1N1 virus in replicate cultures for temporal sampling of influenza-specific CD8 T cells. On days 7, 10, and 14 post-stimulation, all activated T cells (CD3+ HLA-DR+ CD38++) were sorted and analyzed using scRNA-seq and TCR-seq (**Figures 5A, S8A**) to capture antigen-responsive T cells. Transcriptional clustering of the cells revealed 3 dominant profiles of activated CD8 T cells from tonsil organoids: CD8-TRM, CD8-4-1BB+, and proliferating CD8 (**Figures 5B, S8B**). We analyzed the temporal dynamics of these populations and found that both 4-1BB+ and proliferating CD8 populations expanded relative to unstimulated controls (**Figure 5C**).

**Figure 5:**
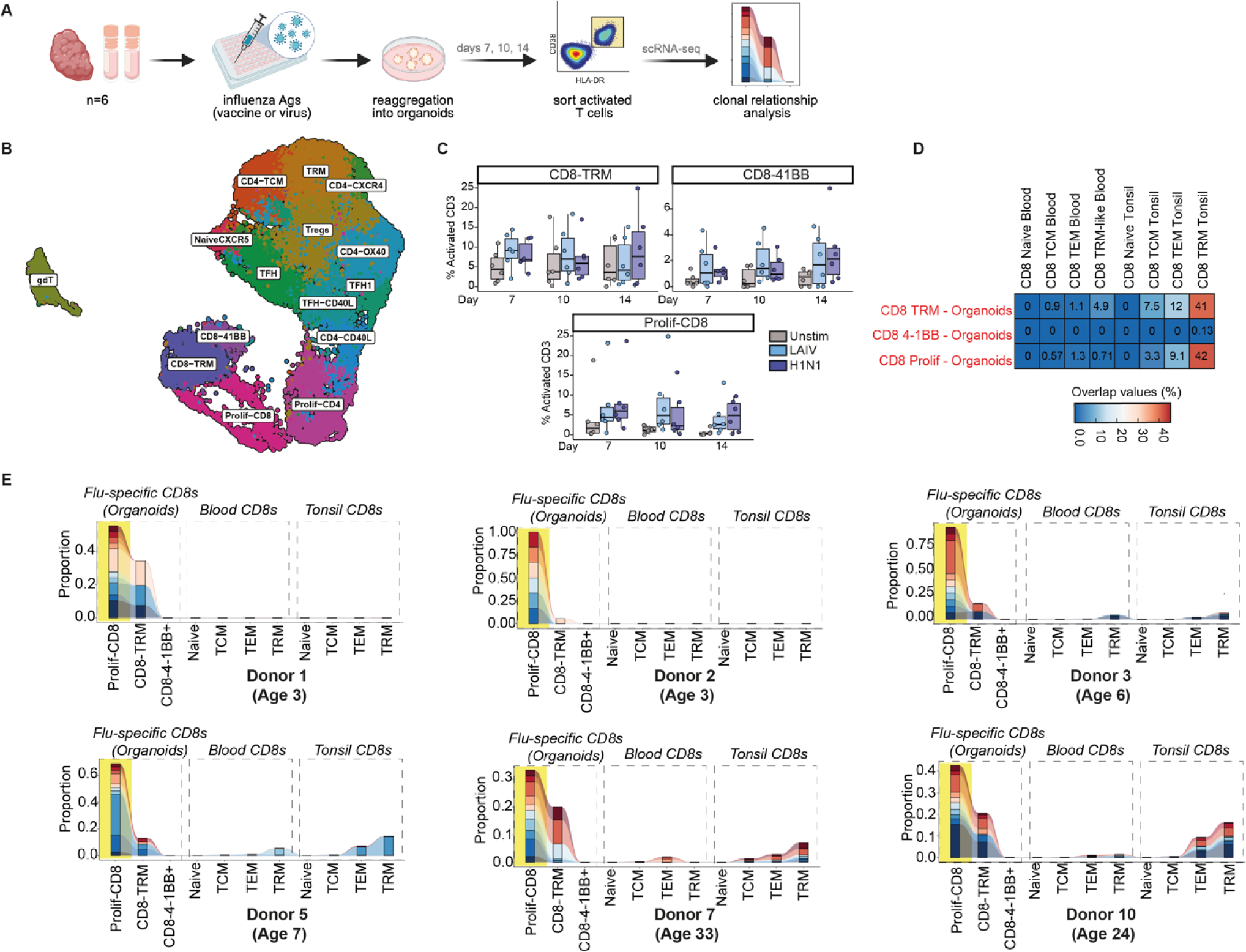
Clonal origins of antigen-specific CD8 T cells following vaccination/infection. (A) Experimental design - tonsil immune organoids were generated from 6 of the 10 donors from the primary cohort and stimulated with live-attenuated influenza vaccine (LAIV) or wild-type virus H1N1. Organoids were harvested on days 7, 10, and 14, and activated CD3+ T cells were analyzed using scRNA-seq. (B) UMAP of activated T cells and their associated annotations based on gene and protein expression. (C) Box plots compare the proportions of CD8 T cell subsets within the pool of total activated T cells over time across stimulation conditions. (D) Heatmap of clonal overlap of flu-specific activated CD8 T cells sampled (x-axis) on days 7, 10, and 14 compared to CD8 T cell subsets from blood and tonsils sampled directly ex vivo (y-axis). The data shown here is aggregated across donors. (E) Clonal tracking analysis of top 10 flu-specific proliferating CD8 T cell clones within each of the 6 donors and their occupied repertoire within CD8 subsets sampled ex vivo from blood and tonsils.

We then compared the clonal relationships between T cells from organoids with CD8 T cell subsets in direct *ex vivo* blood and tonsils from the same individual. Notably, 4-1BB expressing cells were clonally unrelated to other activated CD8 T cells from organoids or any CD8 T cell subsets assessed *ex vivo*, suggesting they differentiated from the naive compartment (**Figures 5D, S8C**). In contrast, proliferating CD8 T cells from antigen-stimulated organoids exhibited significant clonal sharing with CD8 TRM and, to a lesser extent, with CD8 TEM and TCM phenotypes from tonsils (**Figure 5D**). Influenza-specific expanded clones from organoids were under-represented within blood CD8 compartments. Since influenza-specific CD8 T cells are readily detected in circulation following vaccination or infection^57^, our findings suggest that under steady-state conditions, the pool of memory CD8 T cells from the tissue that can respond to influenza antigens mostly consists of resident memory cells.

Granular clonal tracking revealed donor-specific heterogeneity in the relationship between starting and acquired phenotypes after stimulation, presumably influenced by their previous antigen exposure. Organoids from pediatric donors (3 years old) showed no clonal sharing among tonsil memory CD8 subsets, while organoids from adult tissues demonstrated the most sharing (**Figure 5E**). Taken together, these findings indicate that antigen-specific CD8 T cells acquire different phenotypes post-vaccination/infection and that their fate is influenced by their naive or memory origin.

### Tissue-specific phenotypic and clonal segregation of antigen-specific T cells

Recent work demonstrated that antigen specificity is strongly linked to CD8 T cell phenotype^54,58–60^. While this association has been well established in peripheral blood T cells, it remains unclear whether a similar relationship spans different bodily sites. If true, one would expect that CD8 T cells with a tissue-restricted transcriptional profile would have unique TCR specificities compared to blood. To address this, we searched for TCR sequence matches between our data and VDJdb, a public repository of TCRs with known specificities^61^ (**Figure S9**). We identified 22 exact matching TCRs (3,195 CD8 T cells) with VDJdb, some of which were expanded clones in our dataset. We further analyzed the expanded clones by tissue site. For one particular specificity (AVFDRSDAK, derived from EBV), one donor had three different expanded clones, which were present at varying frequencies in blood and tonsils but conserved in phenotype (**Figure S9A**). In another case, two clones specific for a different EBV peptide (FLRGRAYGL) were only found in the tonsils and had TRM and TEM phenotypes (**Figure S9B**). Finally, a sequence identified as a dual binder for a CMV and an influenza peptide (CMV-NLVPMVATAV; influenza-GILGFVFTL) in VDJdb was also found in our dataset and represented a single clone found in circulation (**Figure S9C**), with limited representation in tonsils.

Given these observations from database matching, we next asked if compartmentalization of antigen-specific T cell clones or even specificities extend to other antigens. To test this, we sorted and single-cell sequenced CD8 T cells of known specificities using oligo-barcoded tetramers (**Figure 6A**) with 15 different peptides covering naive antigens (melanoma, yellow fever virus vaccine, human immunodeficiency virus) and a variety of common memory exposures such as Epstein-Barr virus (EBV), cytomegalovirus (CMV), influenza, and severe acute respiratory syndrome-related coronavirus 2 (SARS-CoV-2). Matched blood and tonsils from an independent cohort of donors (2 pediatric and 2 adults) were used for this analysis. Dimensionality reduction and clustering of cells with single antigen specificities revealed clear separation of antigen-specific CD8 T cells by tissue source (**Figures 6B, S10A**). Additionally, these clusters were consistently represented among all donors (**Figure S10B**). While naive and TCM cells were observed in both compartments, TRMs expressing CD69 and CD103 were only found in tonsils, and CD95+ and effector memory RA (EMRA) phenotypes were restricted to blood (**Figures 6B, 6C, and S10A**). We analyzed the tissue compartmentalization and clonality within each antigen specificity and found that clone sizes varied substantially between tissue compartments. EBV and influenza-specific CD8 T cells were more clonal in tonsils than in blood, while CMV-specific cells were clonally expanded in circulation (**Figures 6D, 6E**). Clonal tracking analysis of the top ten clones for each specificity revealed the extent of clonal sharing between compartments (**Figure 6F**). While the top EBV-BMLF clones in either blood (left) or tonsil (right) had representation in the opposing compartment, the top clones specific for EBV EBNA-3C (**Figure 6F**), CMV-pp65, influenza-matrix, and SARS-CoV-2 NP/Spike (**Figure S10B**) were unique within each compartment. These data, combined with our VDJdb matching analysis, demonstrate that most memory TCR clones are compartmentalized to different bodily sites and that this is due, at least in part, to their specificity and the influence of the tissue microenvironment.

**Figure 6:**
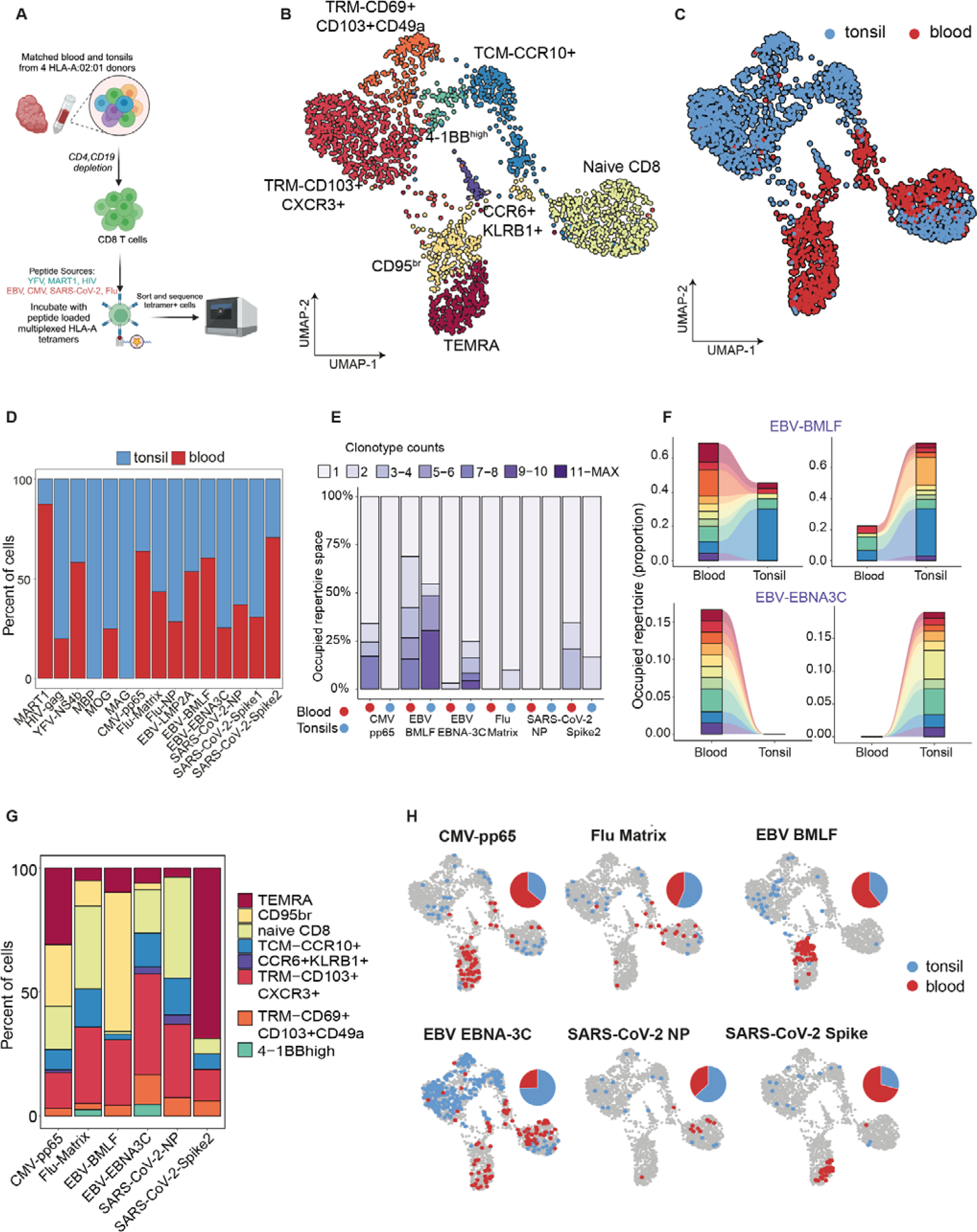
Clonal and phenotypic compartmentalization of Ag+ CD8 T cells. (A) Experimental design for multiplexed assessment of tetramer-specific CD8 T cells from an independent cohort of HLA-A*02:01 donors with tonsils and autologous blood samples. CD4 T cells and B cells were depleted from tonsils and PBMC, stained with multiplexed tetramer assemblies, sorted, pooled, and analyzed using scRNA-seq. (B) UMAP of Ag+ CD8 T cells with associated cluster annotations based on gene and protein markers. (C) UMAP of Ag+ CD8 T cells colored by their tissue origin (D) Distribution of CD8 T cells with different specificities (x-axis) in blood and tonsils. (E) Tissue-specific distribution of clone counts of CD8 T cells with specificities against viral peptides. Data shown here are aggregated across all donors. (F) Tracking of top 10 clones specific to EBV-BMLF (top) and EBV-EBNA3C (bottom) in blood (left) and tonsils (right) and their occupied repertoire in the other tissue compartment. (G) Stacked bar graph comparing the phenotypic distribution of CD8 T cells with specificities against viral peptides. (H) UMAP of CD8 T cells (aggregated across all donors) against memory antigens in blood and tonsils. Individual pie charts highlight the proportion of cells in blood (red) or tonsils (blue).

In humans, significant clonal expansions within the CD8 T cell repertoire are dominated by exposure to chronic infections such as EBV and CMV^62,63^. Our analysis of antigen-specific T cells against these viruses demonstrates that their frequency, phenotype, and clonality vary considerably between blood and tonsils. While CMV pp65-specific cells were largely restricted to blood and acquired an EMRA-like phenotype (**Figures 6G, 6H**), EBV-specific CD8 T cells showed a more complex behavior that was dependent on both the tissue site and peptide specificity. EBNA-3C-specific CD8 T cells (which bind a peptide from a viral latency factor) were found in both the naive and memory CD8 T cell compartments of the tonsil. In contrast, BMLF-specific CD8 T cells (which bind a peptide from an EBV lytic factor) were more likely to be memory cells in circulation (**Figures 6G, 6H**). A similar phenomenon was also true for SARS-CoV-2 proteins (nucleocapsid vs. spike) (**Figures 6G, 6H**). Taken together, these observations suggest that factors such as the site of infection, protein accessibility, and immune evasion mechanisms likely influence the prevalence and differentiation profiles of antigen-specific T cells.

## DISCUSSION

A major impetus behind this study was to define the key determinants of TCR biology by deeply sampling T cells from both blood and tonsils. By comparing these two sources, we aimed to ground our findings with prior research on the peripheral blood TCR repertoire and to contrast them with T cells residing in a mixed lymphoid and mucosal tissue environment. This unprecedented sampling depth analyzed hundreds of thousands to nearly a million single cells from each donor. We juxtaposed this atlas of 5.7 million T cells with cell surface protein readouts from a subset of cells to confidently assign phenotypes widely accepted by the immunology community^64^. Our study lays the groundwork for a definitive understanding of TCR diversity in healthy individuals across various ages and bodily compartments. This foundational knowledge will be a benchmark for identifying disturbances in T cell phenotypes and TCR repertoires related to various human diseases. More importantly, it will aid in the development of advanced analysis tools aimed to quantify and interpret TCR repertoires and reveal complex relationships between TCR sequence, specificity, cell fate, and their respective locations and functions.

### Tissue compartment differences

We hypothesized that T cell phenotypes would differ between tissue compartments. Consistent with our expectations, certain phenotypes, such as TRMs and TFH, were restricted to tissue and rarely detected in circulation during homeostasis. Other memory subtypes, including TCM, TEM, MAITs, and Tregs^56^, were observed in both compartments in varying proportions, sometimes influenced by the donor’s age. Surprisingly, MAITs were more abundant in blood than in tonsils, which likely reflects their rarity in human palatine tonsils rather than discrepancies in cell annotation. Within phenotypes shared between compartments, we observed differences in clonal distributions based on compartmental origin. For example, TCM subsets were more clonal in tissues, while TEMs and MAITs were more clonal in blood, highlighting location-dependent expansion and distribution patterns within phenotypically related cells^65^. While tonsils are just one of the many possible tissue sites, clonal expansions will probably differ based on the tissue and extent of antigen exposure.

Consistent with prior studies, CD8 TEMs were the most clonal among circulating T cell subsets^54^. In tonsils, the largest TCR clones were TFH in the youngest donors and CD8 TEM/TRM in older children and adults. These expanded TFH clones in pediatric donors were transcriptionally diverse and spanned various TFH phenotypes, whereas CD8 TEM clones in adults were transcriptionally homogeneous. This aligns with recent studies showing that clonally related CD8 T cells are more likely to be transcriptionally similar^54^. In contrast, TFH are highly plastic and can co-opt various flavors of Th programs (TFH1, TFH2, TFH17) depending on the immunological context and extracellular cues they sense^66^. We posit this as a possible explanation for the phenotypic diversity of TFH clones.

### Clonal sharing and antigen-specific clones

A major strength of the study was the in-depth sampling of hundreds of thousands of T cells across autologous blood and tissue. While previous studies have investigated the clonality of specific human T cell subsets (TRM, TEM, iNKT) across tissue compartments in humans on a much smaller scale^47,65,67^, the unparalleled sampling in our study (100X deeper on a per donor basis) allowed us to truly understand the extent of clonal sharing and compartmentalization of T cells between blood and tissue. Even at this sampling depth, a key finding from our analyses was low clonal sharing between the compartments, underscoring a consistent theme in patterns of clonal and phenotypic segregation between blood and tissue. The fraction of shared clones was much lower than theoretical estimates of a well-mixed TCR distribution, ranging from 1-7%. Higher levels of clonal sharing were observed in older donors and correlated with increased clonality of memory CD4 and CD8 subsets from tonsils. Another intriguing finding from our analyses was the presence of a population of naive CD4 T cells with small but detectable clonal expansions in both blood and tonsils from adult donors. A similar population has been described for CD8 T cells in older adults^23,68^. While the mechanisms regulating this clonal expansion are unclear^34,69^, we demonstrate that a subset of these cells are clonally related to memory CD4 T cells in tissue. Together, these findings reflect the overall impact of accumulated memory and continuous immunological interactions between blood and tissue with age^25^. Given the differential distribution of T cell phenotypes and clones by location, it is crucial we incorporate additional tissue analyses in human T cell aging studies.

An important discovery from our experiments was the distinct clonality and phenotypic patterns of antigen-specific T cells between blood and tissue. CMV-specific CD8 T cells were often restricted to blood, whereas EBV-specific clones were distributed in blood, tonsils, or both, depending on their specificity. Interestingly, large EBNA3C-specific (latency factor) CD8 T cell clones were confined to tonsils and were undetectable in blood, whereas BMLF-specific (lytic factor) T cells were more likely to be present in circulation. Furthermore, using an *in vitro* organoid model, we demonstrated that influenza-specific CD8 T cells that expanded in stimulated cultures were under-represented in autologous blood repertoire^47,65^. Our analysis shows that localized expansions contribute to disparities in clonal representation, likely dictated by viral presence/function, and suggests that tissue location may play a critical role in establishing immunodominance. It will be crucial to understand the relationship between immunodominant T cell responses and disease or therapeutic outcomes, particularly as they might be influenced by the tissue microenvironment.

One of the key insights from our analysis is that while migration and tissue patrolling are vital T cell functions, this movement is insufficient to establish a full mixing of T cell clones across different bodily sites. Some evidence for this comes from parabiosis experiments in mice^4,70,71^, where tissue compartmentalization was a major contributing factor for immune protection in skin, gut, and lungs during infection. Furthermore, while early activated T cells can migrate to all non-lymphoid organs, memory cell trafficking becomes more restricted over time^72^. The extent of this “division of labor” between circulating and bona fide tissue resident subsets in humans is only beginning to emerge^47,73^. The dynamic nature of T cell trafficking, influenced by factors such as infection site, stage of infection, inflammation, and homeostasis, poses challenges in capturing a consistent picture of T cell distribution and clonal diversity based solely on blood samples^47,65^.

### A renewed estimate of TCR repertoire

The size and complexity of the human immune repertoire have significant implications for areas such as vaccine immunotherapy design, autoimmunity, and immune aging. With increasing age, memory CD8 T cells were more clonal in tonsils than blood, suggesting higher rates of clonal expansion in tissue relative to circulation. In contrast, the CD4 T cell memory subset was consistently more clonal in tissue in all donors. These observations suggest that, in addition to qualitative differences between tissue and blood repertoire, there exists significant differences in clonal distribution between blood and tissue. Current estimates of diversity are derivatives of blood sampling, and range from 10^6^-10^8^, depending on beta vs. paired chain sequencing ^23,33,43^. Our analyses suggest that, at least for the memory compartment, the addition of tissue repertoire provided a better estimate of diversity than blood alone. Furthermore, since blood represents only 2% of total T cells in the human body^3^, we argue that using the tissue repertoire to model the remaining 98% of T cells in the human body would provide a more accurate estimate of TCR diversity. Towards this, our calculations indicate an average total repertoire size of 3.3 × 10^11^ clones based on real T cell distribution in the bodily sites vs. 3.95 × 10^11^ clones based on extrapolation of blood repertoire. This discrepancy is more prominent in young children, with overestimations reaching as high as 42% when only blood was used to predict the overall diversity. Collectively, these findings reiterate that in addition to estimating diversity on a per phenotype basis (naive vs. memory), the source of T cells (blood vs. various tissues and the extent of sharing) are essential factors that should be considered while evaluating the total TCR repertoire in humans at steady state. We argue that current methodologies, which rely on sampling a few thousand cells from circulation, might effectively capture ongoing immune responses to vaccination or immunotherapy, but accurately identifying the peak of T cell response remains challenging.

### Relationship between TCR sequence and phenotype

It has long been believed that T cells with a given specificity differentiating from a common naive cell tend to acquire similar transcriptional or functional profiles. With the advent of single-cell technologies, which permit the integration of protein, gene, and TCR readouts from the same cell, there has been a renewed interest in testing a potential deterministic relationship between a cell’s TCR and its phenotype^54^. It has been reported that CD8 T cells for a given specificity have identical transcriptional profiles in circulation^54^. On the contrary, several studies in mouse models have demonstrated that a single naive CD4^52^ or CD8 T cell^16^ can give rise to multiple fates, suggesting significant interclonal and intraclonal functional heterogeneity in T cell responses. The ultimate fate of the cell depends on the quality of early priming^74^, interactions between T cells and antigen-presenting cells (APC)^75^, or subsequent signals during infection^76^. We hypothesized that the tissue microenvironment could potentially shape the phenotype of clonally related cells in humans. Our analysis revealed high concordance between CD4 vs. CD8 lineages and naive vs. memory status across compartments, suggesting some degree of phenotypic consistency. More specifically, CD8 memory subsets and MAIT cells were more likely to retain similar phenotypic features across blood and tonsils. However, we also observed clonally related T cells with phenotypic disparities across compartments, such as CD8 TRMs in tonsils and CD8 TEMs in blood. This highlights the possibility that identical TCR-bearing clones can adopt divergent functionalities based on location.

We observed a higher degree of clonal sharing between compartments for T cell subtypes present in both blood and tonsils, such as Tregs. This is consistent with a recent report indicating that most Treg clones are specific for ubiquitous self-antigens and widely circulate and recirculate among various organs^56^. Surprisingly, our study points to a significant number of non-Treg cells (TEM, TCM) from blood, that are clonally related to tonsillar Tregs^77^. Similar trends were observed in tissue-restricted TFH phenotypes, which showed significant clonal sharing with non-TFH phenotypes (TEM and TCM) in blood. This is a surprising finding, given that influenza-specific TFH cells found transiently in circulation are clonally related to TFH cells but not to non-TFH cells in the tonsils^78^. Although this may be true transiently during acute antigen stimulation, our analysis highlights the possibility that at a steady state, a TCR clone can acquire distinct phenotypes depending on their localization. More importantly, this finding further underscores the adaptability and plasticity of T cells in response to the microenvironment. Consistent with this idea, a recent study reports the phenotypic plasticity of Spike S_167-180_ specific CD4 T cells in circulation following COVID-19 mRNA vaccination. Mudd et al. showed that spike-specific TFH persisted in draining lymph nodes for 200 days post-vaccination but circulating TFH-like cells were detected only transiently in peripheral blood and acquired a Th1 phenotype 3-4 weeks post-vaccination.

The tissue compartment also had a strong influence on the phenotype of antigen-specific CD8 T cells against common viruses (EBV, CMV, SARS-CoV-2) where cells with identical specificities had TEM or TEMRA phenotypes in circulation but a TRM phenotype in tonsils. The phenotypic fate of antigen-specific CD8 T cells following vaccination or infection was also dictated by the phenotype of their precursors. Analysis of tonsil organoids stimulated with influenza vaccines or viruses allowed us to identify a stimulation-specific pool of resting and proliferating TRMs on day 7 post-stimulation that were clonally related to the pre-existing pool of CD8 TRM or TEM in older children and adults but not in younger children. On the other hand, organoid CD8 T cells expressing the activation marker 4-1BB were clonally unrelated to CD8 T cells sampled on day 0, suggesting their differentiation from the naive pool of TCRs. Taken together, these results highlight the complex relationship between TCR, naive or memory status, and phenotypic fate in the context of antigen stimulation in tissues.

### Limitations of the study

Tonsils are tissue sites that combine lymphoid and mucosal characteristics, making them an ideal location to capture immune responses occurring in these distinct but interconnected environments. However, we recognize that our findings may not be completely generalizable to other tissue sites such as the gut, lung, and skin. Despite the large number of cells studied from a few donors, the observed patterns may not apply to older adult donors, where major age-associated changes in TCR repertoire occur^68,79^. T cell responses are dynamic, and the extent of clonal sharing between tissues might likely be underestimated, particularly during an ongoing infection. Additionally, phenotypic convergence of antigen-specific T cells between blood and tissue during infection is a likely possibility, which was not tested in this study. Finally, although the organoid system aims to capture this dynamic response, microenvironmental factors such as local cytokines and tissue-specific interactions might not be fully understood or easily replicable *in vitro*. Despite these limitations, in toto, this study underscores the limited sharing and selective compartmentalization of T cells and antigen-specific T cell responses, with implications for immune surveillance and tissue-specific immunity. Our study highlights the importance of considering tissue localization when assessing T cell responses, whether for chronic infections, immunotherapy, or vaccination and reinforces the need for paired analysis of blood and affected tissues, when possible, especially for organ-restricted infections.

## RESOURCE AVAILABILITY

### Lead contact

Requests for further information or access to data should be directed to Lisa E. Wagar.

### Materials availability

This study did not generate new unique reagents.

## ACKNOWLEDGEMENTS

The authors thank the tonsillectomy patients and their families for participating in this study, Drs. Jennifer Atwood and Michael Hou at the University of California, Irvine Institute for Immunology flow cytometry core for technical assistance with flow sorting, Drs. Robert Edwards and Delia Tifrea at the University of California, Irvine Medical Center pathology for sample coordination, UCI Genomics Research and Technology Hub (GRT-Hub) for assistance with library preparation, and Drs. Eric Pearlman, Shivashankar Othy, Robert Schreiber, Wayne Yokoyama, Michael Paley, Chyi Hsieh, Ulrike Lorenz, and Maxim Artyomov for critical feedback on the work.

This work is supported by funding from the Wellcome Leap HOPE Program (to L.E.W.) and the National Institutes of Health (NIH) grant R01AI173023 (to L.E.W.). This work was also made possible, in part, by the Genome Technology Access Center at the McDonnell Genome Institute at Washington University School of Medicine for help with genomic analysis. The Center is partially supported by NCI Cancer Center Support Grant P30 CA91842 to the Siteman Cancer Center from the National Center for Research Resources (NCRR), a component of the National Institutes of Health (NIH), and NIH Roadmap for Medical Research. This work is supported by funding from the Children’s Discovery Institute of Washington University and St. Louis Children’s Hospital (MI-LI-2020-914 to N.S.). This work was also made possible, in part, through access to the following: the Genomics Research and Technology Hub (formerly Genomics High-Throughput Facility) Shared Resource of the Cancer Center Support Grant (P30CA-062203), the Single Cell Analysis Core shared resource of Complexity, Cooperation, and Community in Cancer (U54CA217378), the Genomics-Bioinformatics Core of the Skin Biology Resource Based Center (P30AR075047) at the University of California, Irvine and NIH shared instrumentation grants 1S10RR025496-01, 1S10OD010794-01, and 1S10OD021718-01. This publication is solely the responsibility of the authors and does not necessarily represent the official view of the funding agencies.

## AUTHOR CONTRIBUTIONS

Designed, optimized, and performed experiments: SS, EL, JMK, AD, MTM, SJHM, CT

Analyzed and interpreted data: SS, JH, SB, AB, KB, LEW, AT-M, NS

Sample procurement: AMS, DT, GA, QZ

Conceived of study, study design, secured funding: SS, AM, AT-M, NS, LEW

Writing the original manuscript: SS, LEW

Edited, reviewed, and approved manuscript: all authors.

## METHODS

### Informed consent and sample collection

Tonsils from healthy consented individuals undergoing surgery for obstructive sleep apnea, hypertrophy, or recurrent tonsillitis were collected in accordance with the University of California, Irvine Institutional Review Board (IRB). Ethics approval was granted by the University of California, Irvine IRB (protocol #2020-6075), and all participants provided written informed consent. Participants in the primary cohort used for the study were aged 3-39 (**Table S1**). Overall, tonsil tissue was healthy in appearance. Although most donors were otherwise healthy, two of the participants self-reported prior diagnosis with an autoimmune disease (psoriatic arthritis in Donor 8 and rheumatoid arthritis in Donor 10).

### Sample processing

Samples were processed as previously described. Briefly, whole tonsils were collected in saline after surgery and immersed in an antimicrobial bath of Ham’s F12 medium (Gibco) containing Normocin (InvivoGen), penicillin, and streptomycin for 30-60 min at 4°C for decontamination of the tissue. Tonsils were then briefly rinsed with PBS and mechanically dissociated; debris was removed using gradient centrifugation (Lymphoprep, Stemcell). PBMCs were isolated from blood (diluted 1:1 in PBS) by standard density gradient centrifugation over Ficoll. Cells were counted, and samples were cryopreserved in fetal bovine serum (FBS) with 10% DMSO and stored in nitrogen until use.

### Pathogen exposure history

Prior exposure to select viruses was determined using semi-quantitative ELISAs measuring IgG antibodies against CMV (Abcam, cat #ab108724), EBV-VCA (Abcam, cat #ab108730), and EBV-EBNA1 (Abcam, cat #108731) per manufacturer’s instructions. Briefly, diluted serum samples were added to pre-coated wells and incubated at 37’C for 1 hour. Samples were then washed, treated with HRP-conjugated secondary antibodies, washed, and TMB substrate was used for signal development. Experimental readouts were measured colorimetrically and seropositivity was established using the thresholds defined by the manufacturer for each kit.

### T cell isolation for scRNA-seq

Cryopreserved tonsil and PBMC samples (donors 1-10) were revived using pre-warmed media: RPMI1640 supplemented with glutamax, 10% heat-inactivated FBS, 1x nonessential amino acids, 1x sodium pyruvate, and 1x penicillin-streptomycin. Cells were washed then washed with FACS buffer (PBS + 2% FBS + 0.05% Sodium Azide). Roughly 10^7^ PBMC or tonsil cells were stained with Live/Dead NIR (1/1500) in PBS for 30 minutes in the dark. Cells were washed and prestained with Fc Block (1/50) for 20 minutes on ice and stained with a cocktail of antibodies - CD3-BB700 (1/50), TCR-gd-PE-Cy7 (1/100), CD14-BV605 (1/100), CD19-BV605 (1/100), and CD16 (1/100) for an additional 30 minutes on ice. After washing, cell pellets were resuspended in 200 uL of 1X PBS + 0.04% BSA and placed on ice prior to sorting. ab+ T cells were sorted based on viable single cells with markers CD19/CD14/CD16-CD3+ TCRgd-(**Figure S1A**).

### scRNAseq with scTCR

Sorted cells were washed and resuspended in 1X PBS with 0.04% BSA, counted, and resuspended in a final concentration at 1200 cells/uL. Single-cell suspensions were loaded on a Chromium X controller (10X Genomics) with a loading target of 30,000 - 50,000 cells in several replicates (4-24 GEM samples depending on the sort yield) with a goal of sequencing ∼500,000 cells per donor per compartment (blood or tonsils). Gene expression and TCR libraries were generated using the Chromium Next Gem Single Cell 5’ Reagent Kit v2 (Dual Index) per the manufacturer’s instructions. Quality and quantity of libraries were measured on tapestation, qubit, and bioanalyzer and sequenced on Illumina NovaSeq 6000 with a sequencing target of 30000 reads per cell for gene expression libraries and 5000 reads per cell for TCR.

### scRNAseq with cell surface protein analysis

Cryopreserved tonsil and PBMC samples from 8 independent donors (donors 11-18, ages 3-58) (**Table S1**) were revived in pre-warmed RPMI1640 (supplemented with 10% FBS), washed with FACS buffer, and stained using a cocktail containing Fc block, CD19-APC (1/50), CD3-AF488 (1/50), and TCRgd-PE (1/50). Additionally, during the antibody staining step, samples were stained with unique hashing antibodies (TotalSeqC, BioLegend CA) for sample barcoding. Cells were stained for 30 minutes on ice, washed twice with FACS buffer, and sorted on a FACSAria Fusion (BD Biosciences). Roughly 50,000 ab+ T cells were collected from viable single cells with CD19-CD3+TCRgd-phenotype, pooled by compartment, washed with FACS buffer, and stained with a cocktail of oligo-tagged cell surface antibodies (**Table S2**). Cell pellets were washed thoroughly in 4 mL FACS buffer for a total of 4 washes. After the final wash, pellets were resuspended in 100 uL 10X buffer, counted, and resuspended to a final concentration of 2,400 cells/uL. Single-cell suspensions were loaded on a 10X Genomics Chromium controller in duplicates with a loading target of 60,000 cells per sample. Libraries were generated using the Chromium Next Gem Single Cell 5’ Reagent Kit v2 (Dual Index) per the manufacturer’s instructions with the addition of 5’ Feature Barcode libraries. Quality and quantity of libraries were measured on tapestation, qubit, and bioanalyzer, and sequenced on Illumina NovaSeq 6000 with a sequencing target of 30,000 reads per cell for gene expression libraries and 10,000 reads per cell for TCR and feature barcode libraries.

### Single-cell data QC

Raw reads from gene expression, TCR, and for a subset of donors, feature barcode libraries were aligned using Cell Ranger Single Cell Software Suite with Feature Barcode addition (version 6.1.1; 10X Genomics) against the GRCh38 human reference genome (GRCh38-2020-A) using the STAR aligner (version 2.7.2a). Alignment was performed using feature and vdj options (vdj_GRCh38_alts_ensembl-5.0.0) in Cell Ranger. Samples were aggregated, resulting in one composite alignment file for each donor and compartment. For the 8 donors with paired cell surface data, feature files from Cell Ranger were manually updated to separate hashing antibodies (HTO) vs. other cell surface antibodies (Protein) to facilitate independent normalization procedures. Next, data frames from each donor/compartment were split by libraries for QC. For each library, the gene expression matrix was loaded in Seurat (version 5.0.2) and TCR genes were summarized into custom *TCRA*, *TCRB*, *TCRG*, and *TCRD* genes to avoid downstream clustering based on V/D/J gene usage. Cellular features such as mitochondrial and ribosomal gene expression and the number of housekeeping genes were added as additional meta-features. Next, within each library, doublets were identified computationally, with an expected doublet rate of 0.4% per 1,000 cells. DoubletFinder (version 3)^80^ was run on default settings (pN = 0.25, pK = 0.09, PCs=1:20) and removed from subsequent QC steps. Poor quality cells and droplets with ambient RNA (number of expressed genes less than 200 and housekeeping genes less than 25) and additional doublets (number of expressed genes more than 4000) were excluded from subsequent analysis. Finally, libraries from each donor and compartment were re-merged to generate a QC-filtered dataset.

### Donor specific analysis

Before integrating data from all 10 donors, QC-filtered data from blood and tonsils from each donor were analyzed independently. Donor specific objects were generated by merging blood and tonsil datasets from each donor using the *merge* function in Seurat. Data normalization and variance stabilization were performed on the merged donor-specific object using *NormalizeData* and *ScaleData* functions in Seurat. Dimension reduction was performed using the *RunPCA* function to obtain the first 30 principal components, of which the first 20 were used for clustering with Seurat’s *RunUMAP* function with a resolution of 0.8. At this stage, cell types were broadly assigned as *CD3D* expressing T cells and *CD19* or *MS4A1* expressing B cells. Contaminating B cell clusters from each UMAP were filtered before downstream integration. For samples with paired protein information, data were normalized using centered log-ratio transformation (CLR), and donor assignment and doublet removal were performed using the *HTODemux* function in Seurat^81^. For cell surface protein analysis, read counts were normalized using CLR, and any contaminating CD19 protein-expressing clusters were excluded from downstream integration.

### Data Integration and harmonization

To enable efficient and large-scale integrative analysis of millions of cells, we used atomic sketch integration in Seurat^82^. This involved a) iterating through each donor, sampling 50,000 cells (atoms) each from blood and tonsils with equal representation, b) generating a dictionary representation to reconstruct each cell based on the atoms, c) integrating the atoms from each dataset using *FastRPCAIntegration*, and d) for each donor, reconstructing each cell from the integrated atoms using *IntegrateSketchEmbeddings*. Since this approach does not require loading or processing the entire dataset simultaneously, it easily enables analysis of all 5.7 million cells. For the integrated object, we ran *FindVariableFeatures* to extract the top 2,000 variable genes, performed z-score transformation using *ScaleData*, and then performed principal component analysis using *RunPCA*. To correct batch effects, we used Harmony v1.0.107 to scale millions of cells across donors but not compartments. We used *RunHarmony*, with the top 30 PCs as input, and corrected batch effects across donors^83^. The success of data integration and harmonization was tested qualitatively with the *FindClusters* and *RunUMAP* function using the top 15 dimensions and resolution 2.0, resulting in 47 clusters encompassing 5.7 million cells from blood and tonsils combined.

### Cluster annotation

Slight over-clustering of the dataset allowed us to annotate clusters to a finer level (annotation level 4), which were progressively consolidated to broad T cell annotations (levels 3, 2, and 1). Gene and protein markers for each of the 47 clusters were identified using the *FindMarkers* function in Seurat. For gene expression, we used Wilcoxon Rank Sum tests for differential marker detection using a log2 fold change cutoff of at least 0.4, and clusters were identified using a known catalog of markers defined for T cells from human tonsil^64^. A similar approach was followed for protein expression. Both positive and negative protein markers from each cluster were used to aid manual annotation. In certain cases, clusters were merged when no distinguishing markers were identified between cell states (E.g., TFH subsets). A list of the final cluster-specific markers is provided in **Table S3**.

### TCR preprocessing

Only QC-filtered cells with confirmed annotation were included in downstream TCR analysis. In addition, cells were included in the TCR analysis only if they contained a single TRB chain and a maximum of 2 TRA chains as productive TCRs. In a further processing step, only the TRA with the highest UMI was retained for cells with multiple TRA to simplify downstream analyses. A secondary dataset was created to analyze clonal expansion, aggregating cells with the same clonotype. Clonotypes were defined by identical full TRA+TRB nucleotide sequences Clonotypes were annotated by the majority origin (blood or tonsil), and phenotypic annotations of corresponding cells. Basic features of the repertoire (clonality by donor, cluster, compartment) was described using standard workflows in Immunarch^84^.

### Clonal overlap and mixing

Data were filtered for clonal overlap/mixing analysis to retain only cells annotated as memory phenotype. The following filtering steps were performed for clonal mixing/sharing analysis to equalize blood and tonsil clone size distributions. For each donor, only clones with sizes from 10-100 cells were retained. The number of unique clones in blood and tonsils was equalized by subsampling, followed by equalizing the total number of cells in blood and tonsils. Cells from blood or tonsils were randomly permuted among all cells for a given donor to create a well-mixed dataset. The number of counts for a given clone in blood and tonsils was calculated to compute mixing fractions for the true and well-mixed sets. The probability that a member of a given clone group would be found in a blood sample was then computed as:

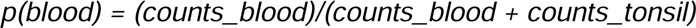

and vice versa. This subsampling was performed 10 times for each donor. The data from each run were aggregated to produce the distribution of mixing.

### Diversity Estimates and Phenotypic matching

T-cell richness was computed using the Chao1 diversity measure^51^. Phenotypic coincidence probabilities were calculated using an unbiased Simpson’s diversity estimator^85^, with conditional coincidence probabilities calculated by an equally weighted average across conditioning groups. Sampling variance was calculated using the unbiased variance estimator described in^86^. Rarefaction curves were generated by repeated random subsampling of cells and calculating the number of clones within each sample. For each donor, these curves were extrapolated to total T cells in the body, based on estimates published in^3^ solely from tonsils or blood repertoire or a combination of both (98% tonsil and 2% blood).

### VDJ-db queries

A matching function was used to query our dataset and associated metadata (donor-specific TCRs, HLA, and annotations) against a set of reference TCRs with known specificities from VDJ-db (cite). Intersecting matches were based on common features such as TRB amino acid sequences and paired TRA-TRB amino acid sequences. Once matched, the epitope and epitope species from the reference and a list of matching cell barcodes were retrieved. Matching sequences were further filtered to ensure either class-I or class-II allele matching for a particular donor.

### Immune organoids for analysis of activated CD8 T cells

Cryopreserved tonsil cells from 6 of the original 10 donors (donors 1, 2, 4, 5, 7, and 10) were thawed, enumerated, and roughly 1.5 e6 cells were plated in 200 µl final volume in ultra-low attachment plates (Corning). Organoid media was composed of RPMI1640 with glutamax, 10% FBS, 1x nonessential amino acids, 1x sodium pyruvate, 1x penicillin-streptomycin, 1x Normocin (InvivoGen), 1x insulin, selenium, transferrin supplement (Gibco), and 0.5 µg/ml recombinant human BAFF. The following amounts of influenza antigens were added to replicate cultures on day 0 based on our prior titration^87^: A/California/07/2009 H1N1 virus - 2.5 hemagglutination units (HAU) per culture; 2019/20 live attenuated influenza vaccine (LAIV FluMist® Quadrivalent) - 1/2,000 final dilution. Cultures were incubated at 37°C, 5% CO_2_ with humidity, and media was replenished every other day by exchanging 30% of the volume with fresh organoid media. Organoids were cultured for two weeks and harvested on days 7, 10, and 14. On harvest days, organoids were washed in FACS buffer and surface stained in a cocktail containing CD19-BV605 (1/100), CD3-APC-Cy7 (1/100), HLA-DR-PerCP-Cy5.5 (1/100), CD38-FITC (1/100), and Zombie Aqua (1/200), in addition oligo-tagged surface antibodies (TotalSeq-C, **Table S2**). Additionally, each organoid was labeled with a unique hashing antibody (TotalSeq-C), allowing for sample multiplexing. Samples were stained for 30 minutes in the dark on ice, washed twice with FACS buffer, and left on ice until sorting. Activated T cells were sorted from viable singlets from the CD3+HLA-DR++CD38++ gate. Cells were washed with 10X buffer, counted, resuspended in a final concentration of 2,400 cells/uL, and immediately loaded into a 10X Genomics Chromium Controller. Libraries were generated using the Chromium Next Gem Single Cell 5’ Reagent Kit v2 (Dual Index) per the manufacturer’s instructions with the addition of TCR and 5’ Feature Barcode libraries. Quality and quantity of libraries were measured on tapestation, qubit, and Bioanalyzer and sequenced on Illumina NovaSeq 6000 with a sequencing target of 30,000 reads per cell for gene expression libraries and 10,000 reads per cell for TCR and feature barcode libraries.

### scRNA analysis of activated CD8 T cells

Sequencing reads were aligned using Cell Ranger, with feature barcode and hashing antibody readouts. Data QC was performed in Seurat using cutoffs described above. Additional doublets were removed based on co-expression of multiple hashing antibodies. QC filtered objects from day 7, 10, and 14 libraries were merged in Seurat. For the integrated object, we ran *FindVariableFeatures* to extract the top 10,000 variable genes, performed z-score transformation using *ScaleData*, and then performed principal component analysis using *RunPCA*. Finally, we ran *FindClusters* and *RunUMAP* function using the top 20 dimensions and resolution 0.8, resulting in 15 clusters of activated T cells encompassing 26,723 cells from tonsil organoids. Clusters were annotated based on protein readouts and top gene markers calculated using *FindAllMarkers* in Seurat. Three CD8 T cell subsets were further analyzed for clonal sharing and tracking with *ex vivo* CD8 T cell subsets from donor-matched blood and tonsils using Immunarch^84^.

### Antigen-specific CD8 T cell analysis

Antigen-specific CD8 T cells were profiled using Chromium Single Cell 5’s Barcode Enabled Antigen Mapping (BEAM) technology (10X Genomics). Peptides (Genscript, 100 uM each) from commonly circulating viruses (EBV, CMV, influenza, SARS-CoV-2) or sources naive to most individuals (myelin, melanoma, YFV, HIV) were prescreened on an HLA-A*02:01 donor per the manufacturer’s instructions and analyzed by flow cytometry. On experiment day, assemblies were generated for 15 peptides (**Table S4**), and a negative control and incubated overnight at 4C. The following day, each of the assemblies (negative controls and target peptides) was quenched for 15 minutes on ice per the manufacturer’s instructions, pooled, and left on ice in the dark. CD8 T cells from autologous blood and tonsils from 4 HLA-A*02:01 donors were enriched using negative selection magnetic beads (CD4, CD19 depletion, Miltenyi Biotec). Roughly 1.5 e6 enriched CD8s were stained with Fc block for 10 minutes on ice, mixed with pooled BEAM assemblies, and incubated for an additional 15 minutes on ice in the dark. Samples were stained with a cocktail containing Zombie Aqua (1/200), CD3-BV605 (1/50), CD19-APC (1/50), CD4-Pacific Blue (1/50), CD8-APC-Cy7 (1/100), CD45RA (1/50) prepared in FACS buffer. The staining cocktail also contained a universal cocktail of oligo-tagged antibodies (**Table S2**) and unique hashing antibodies (BioLegend) for sample barcoding. Samples were incubated for 30 minutes on ice in the dark, washed twice with FACS buffer, and sorted on FACSAria Fusion (BD Biosciences). CD19-CD3+CD4-CD8+ PE+ cells were sorted into 10X buffer, washed twice, resuspended in 25 uL, and loaded on the 10X Genomics Chromium Controller. Libraries were generated using the Chromium Next Gem Single Cell 5’ Reagent Kit v2 (Dual Index) per the manufacturer’s instructions with the addition of TCR, BEAM, and 5’ Feature Barcode libraries. Quality and quantity of libraries were measured on tapestation, qubit, and Bioanalyzer and sequenced on Illumina NovaSeq 6000 with a sequencing target of 30,000 reads per cell for gene expression libraries and 10,000 reads per cell for TCR, BEAM, and feature barcode libraries. Data were aligned and analyzed as described for activated T cells, but with the incorporation of BEAM libraries in Cell Ranger multi alignment, which provided antigen-specificity scores for each cell after correcting for signals from negative control peptide. Only cells with a single dominant peptide specificity score (cutoff 0.5) were included in downstream analyses.

## SUPPLEMENTAL INFORMATION

Figure S1: T cell profiles from blood and tonsils.

Figure S2: Donor heterogeneity in T cell profiles.

Figure S3: Phenotypes of largest T cell clones in individual donors.

Figure S4: Clonal mixing between blood and tonsils.

Figure S5: Distribution of shared clones between blood and tonsils.

Figure S6: Diversity estimates of human blood and tissue TCR repertoire.

Figure S7: Granular annotations of T cell clusters.

Figure S8: Clonal origins of CD8 T cells responding to influenza vaccination/infection.

Figure S9: Predicted specificities of expanded CD8 T cell clones.

Figure S10: Phenotype and clonal distribution of Ag+ CD8 T cells in blood and tonsils.

Table S1: Donor characteristics and exposure history.

Table S2: List of TotalSeq-C antibodies used in this study.

Table S3: Top gene and protein markers across clusters at annotation level 4.

Table S4: List of peptides used in multiplexed class I tetramer experiment.

